# Mode of action of the toxic proline mimic azetidine 2-carboxylic acid in plants

**DOI:** 10.1101/2024.06.10.598327

**Authors:** William Thives Santos, Varun Dwivedi, Ha Ngoc Duong, Madison Miederhoff, Kathryn Vanden Hoek, Ruthie Angelovici, Craig A. Schenck

## Abstract

Plants have an amazing capacity to outcompete neighboring organisms for space and resources. Toxic metabolites are major players in these interactions, which can have a broad range of effectiveness by targeting conserved molecular mechanisms, such as protein biosynthesis. However, lack of knowledge about defensive metabolite pathways, their modes of action, and resistance mechanisms limits our ability to manipulate these pathways for enhanced crop resilience. Nonproteogenic amino acids (NPAAs) are a structurally diverse class of metabolites with a variety of functions but are typically not incorporated during protein biosynthesis. Here, we investigate the mode of action of the NPAA azetidine-2-carboxylic acid (Aze), an analog of the amino acid proline (Pro). Using a combination of plate-based assays, metabolite feeding, metabolomics, and proteomics, we show that Aze inhibits the root growth of Arabidopsis and other plants. Aze-induced growth reduction was restored by supplementing L-, but not D-Pro, and non-targeted proteomics confirms that Aze is misincorporated for Pro during protein biosynthesis, specifically on cytosolically translated proteins. qRT-PCR analysis and free amino acid profiling show that the unfolded protein response is upregulated during Aze treatment implicating protein degradation of misfolded proteins. This study demonstrates the mode of action of Aze in plants and provides a foundation for engineering Aze production and tolerance in crops for enhanced resilience.

## Introduction

Plants have evolved a structurally diverse repertoire of metabolites to enable interaction with the surrounding environment. Thus, plant metabolites have many functions including attracting pollinators and seed dispersers, recruitment of beneficial microbes, deterring feeding insects and surrounding organisms, competition for space and resources, and mitigation of abiotic stresses (Erb and Kliebenstein, 2020; Weng et al., 2021; Bai et al., 2024). Most of these metabolites fall within the category of specialized metabolites, which are lineage and tissue specific metabolites that are biologically active and often more structurally complex than their precursors from core metabolism (Moghe and Last, 2015; Schenck and Last, 2020). As a group, plants make on the order of one million unique and structurally complex metabolites (Rai et al., 2017; Alseekh and Fernie, 2018).

Nonproteogenic amino acids (NPAAs) are a group of plant metabolites with diverse structures and functions (Huang et al., 2011). NPAAs are structurally defined as an amino acid (amino and carboxylic acid groups bound to an alpha-carbon) but are not typically used in protein biosynthesis (Bell, 1976) unlike the 20 common proteogenic amino acids. Some NPAAs are widely distributed across plants and act as intermediates in core metabolism such as S-adenosylmethionine and ornithine (Bell, 2003; Huang et al., 2011; Vranova et al., 2011). Whereas other NPAAs are restricted to certain clades and serve roles in defense, such as meta-tyrosine, canavanine, and mimosine (Brewbaker and Hylin, 1965; Rosenthal, 1990; Bertin et al., 2007; Huang et al., 2012). The structures and distribution of numerous NPAAs across plants have been defined yet, their biological activities and modes of action remain poorly understood.

Azetidine 2-carboxylic acid (Aze) is a NPAA that is produced in phylogenetically distinct plants including *Convallaria majalis* (lily of the valley) and *Delonix regia* from the Asparagales and Fabales orders, respectively (Fowden and Steward, 1957; Sung and Fowden, 1971; Leete et al., 1974; Minakata et al., 1985) as well as from table beet (Caryophyllales; Rubenstein et al., 2006). Aze is a structural analog of the amino acid proline (Pro, Fig. 1A), and inhibits the growth of plants, bacteria, and insects (Adeyeyé and Blum, 1989; Lee et al., 2016; Biratsi et al., 2021). Aze disrupts protein folding as demonstrated by the aggregation of collagen a proline rich protein when Human skin fibroblasts were grown on media containing Aze (Tan et al., 1983). Aze toxicity has also been investigated in plants; Arabidopsis roots and callus cultures were previously shown to be sensitive to growth on Aze (Lee et al., 2016). Arabidopsis has two Pro-tRNA synthetases, enzymes which activate amino acids and deliver them to the ribosome for protein synthesis (Rubio Gomez and Ibba, 2020). One Arabidopsis Pro-tRNA synthetase is specific for Pro over Aze, but the other uses Pro and Aze equally (Lee et al., 2016). A highly specific Pro-tRNA synthetase enzyme that selectively uses Pro and excludes Aze was identified in the Aze-producer *D. regia* (Norris and Fowden, 1972). These studies suggest a role for Aze in the disruption of protein biosynthesis, yet the mode of action in plants remains unexplored.

**Fig. 1.**
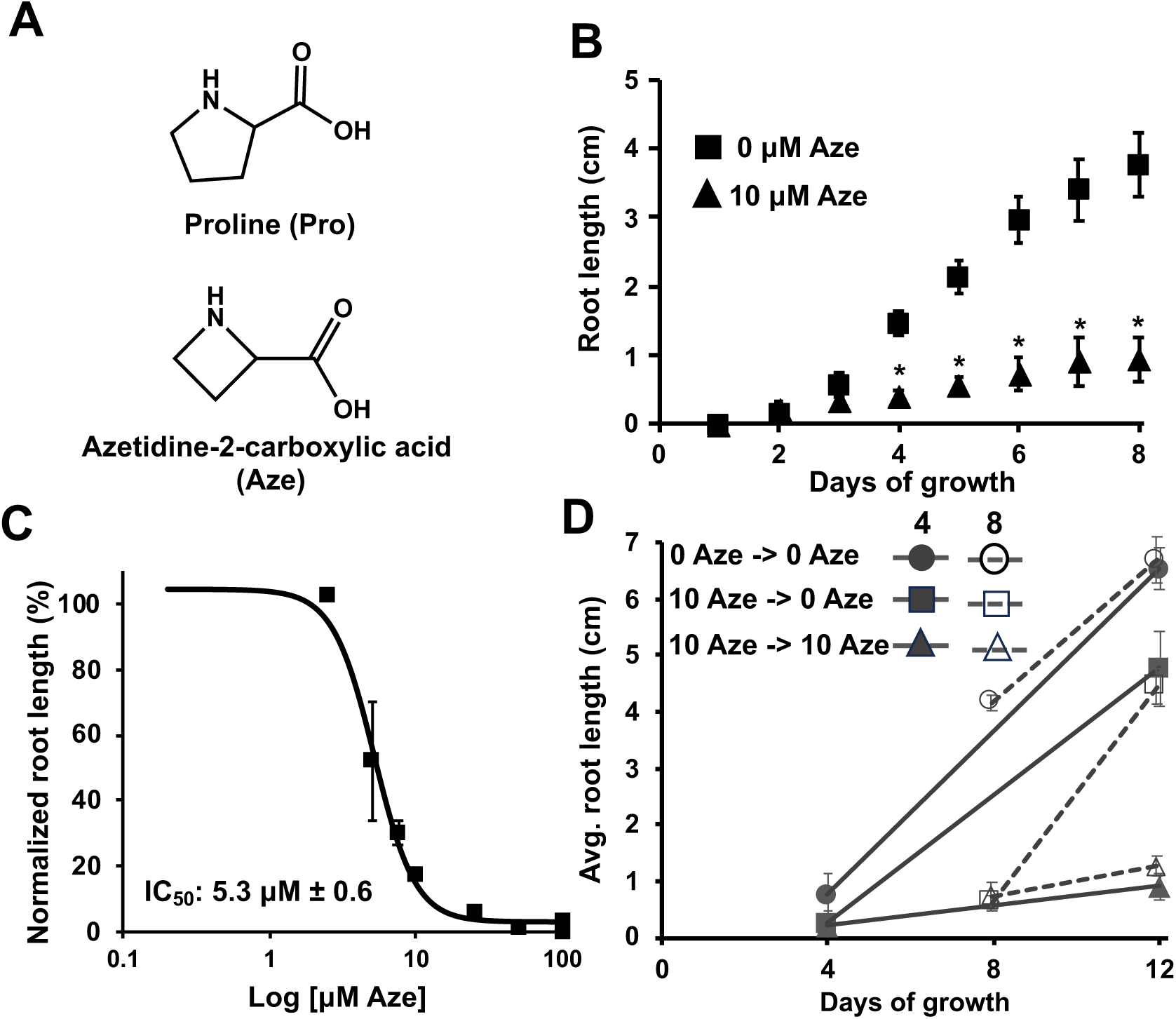
Arabidopsis response to growth on Aze and reversibility. **(A)** Structures of Aze and Pro **(B)** Arabidopsis was grown on 0 (boxes) or 10 μM Aze (triangles) and root length was monitored every 24 hours. Points indicate mean ± SEM of ≥ 8 biological replicates. Stars indicate significant differences between Aze and control *P* < 0.05 in unpaired *t*-test. **(C)** Arabidopsis was grown on increasing Aze concentrations from 2.5-100 μM. Symbols represent mean ± SEM of ≥ 7 biological replicates. **(D)** Arabidopsis was grown on 10 μM Aze for 4 (filled symbols) and 8 (open symbols) days and transferred to media with (open triangles) and without (open circles and squares) Aze. Points indicate mean ± SEM of ≥ 8 biological replicates.

Here we investigate the mode of action of Aze in Arabidopsis and other plants using a combination of plate-based assays, amino acid profiling, and proteomics. We find that Aze is a broad inhibitor of plant growth that can be reversed by supplementation with L-Pro, but not D-Pro. Aze is misincorporated during protein biosynthesis in Arabidopsis specifically on nuclear encoded genes translated in the cytosol and triggers the unfolded protein response (UPR). These results demonstrate the mode of action of Aze in plants and provide the foundation for understanding how plants can produce Aze and avoid autotoxicity.

## Results

### Diverse plants are inhibited by Aze

To determine the biological effect of Aze on plant growth, Arabidopsis was grown on plates supplemented with 10 μM Aze. Growth was monitored every 24 hours for an eight-day period. In the absence of Aze, Arabidopsis displays a near linear growth curve following germination at about 2 days (Fig. 1B). In contrast, when 10 μM Aze was added into the media, a linear growth curve was observed, yet at a much slower rate (Fig. 1B). Germination rate was not affected when Arabidopsis was grown on 10 μM Aze. To characterize the effective concentration range of Aze, Arabidopsis was grown on media supplemented with varying concentrations of Aze, from 2.5 to 100 μM. Complete lack of growth and impacts on germination were observed at 100 μM (Fig. 1C). This concentration gradient allowed us to calculate an IC_50_ for Aze of 5.3 μM ± 0.6 (Fig. 1C). To test if Arabidopsis growth could be restored following Aze treatment, Arabidopsis was grown on 10 μM Aze for 4 or 8 days, then transferred to control media and growth was monitored for a total of 12 days (Fig. 1D). After both 4 and 8 days of 10 μM Aze treatment, growth could be fully restored, suggesting that growth reduction can be overcome after removing Aze from the media.

Next, we tested if a broader range of plants are inhibited by the presence of Aze. We tested model and non-model plants as well as some crops including lettuce, tobacco, tomato, *Nicotiana benthamiana*, and *Psychine stylosa*. We performed a root growth assay on plates in the presence of various Aze concentrations from 2.5 – 80 μM. Root growth was inhibited in all the plants that were tested, however the amount of Aze required to reduce root growth varied (Supplementary Fig. 1). For example, *P. stylosa* (within the Brassicaceae family) was very sensitive to Aze and 90% root reduction was observed at 5 μM Aze, whereas lettuce was more tolerant and 40 μM Aze was required to reduce root growth by 90% (Supplementary Fig. 1).

These data suggest that Aze is broadly toxic to plants and its effect on plant growth varies based on the system and concentration of Aze.

### Restoration of Aze growth reduction by supplementation with Pro

To test whether Arabidopsis growth on Aze could be restored by Pro supplementation, plants were grown on 10 μM Aze plus 100 μM of L-Pro (Fig. 2A). Root growth was restored completely by Pro supplementation (Fig. 2A). Next, we grew Arabidopsis on 10 μM Aze and varied L-Pro concentrations from 10-200 μM. Root inhibition of 80% was observed in the presence of 10 μM Aze (Fig. 2A,B). When Aze and Pro were provided in a 1:1 molar ratio (10 μM each of Pro and Aze) growth was restored but only slightly (Fig. 2B). Full growth restoration was observed at an 8:1 Pro:Aze ratio (80 μM Pro and 10 μM of Aze) (Fig. 2B) similar to previous results (Lee et al., 2016). To test if the stereoisomer D-Pro could restore growth, 100 μM D-Pro was supplemented to 10 μM Aze media, however this had no effect on restoring root growth (Fig. 2A).

**Fig. 2.**
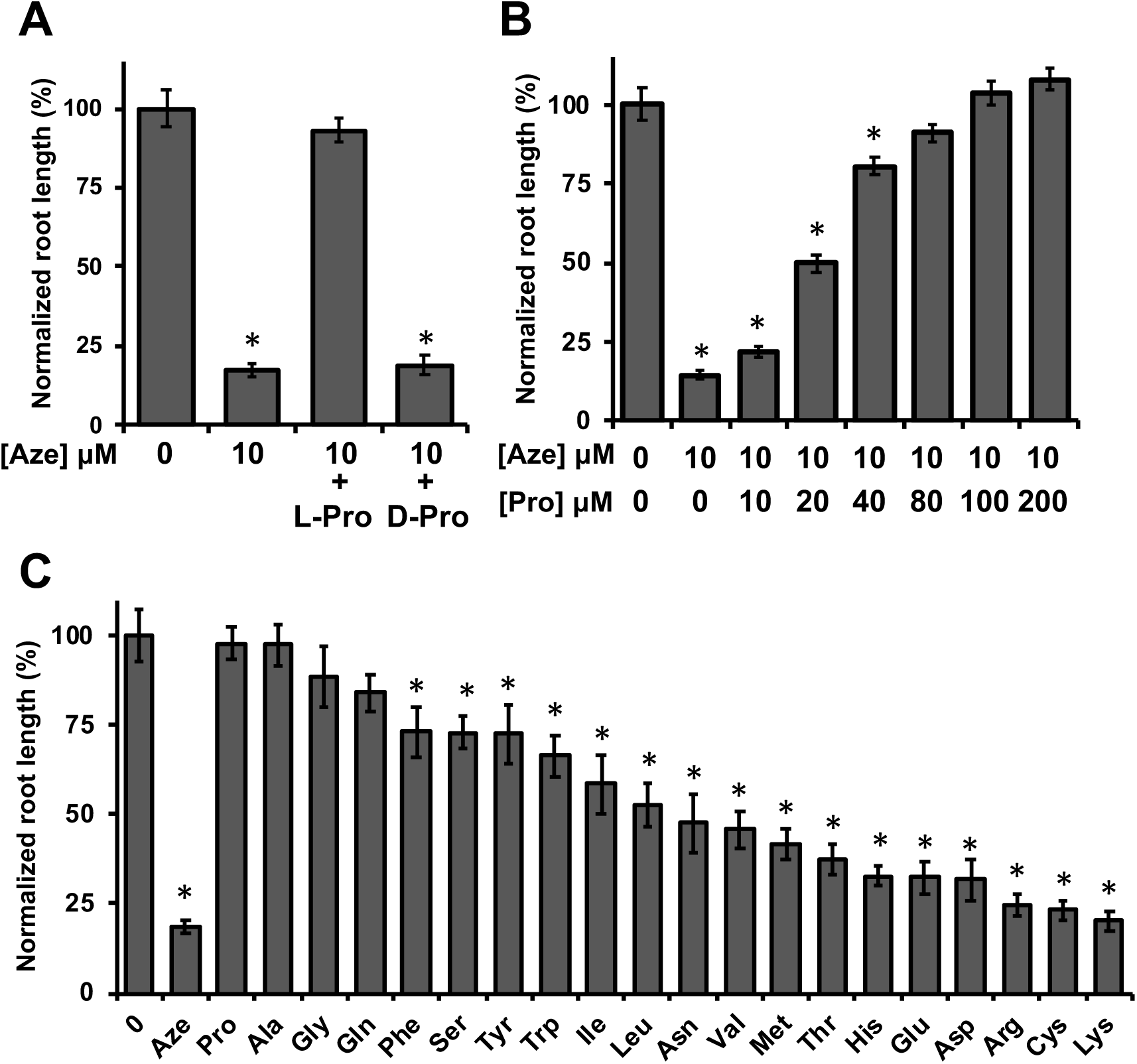
Recovery of growth on Aze with Pro and other amino acids. **(A)** Arabidopsis was grown on 10 μM Aze plus addition of 100 μM of L- or D-Pro. **(B)** Arabidopsis was grown on 10 μM Aze plus varying concentrations of L-Pro. **(C)** Arabidopsis was grown on 10 μM Aze plus addition of 100 μM of the other 19 proteogenic amino acids. All amino acids were the L stereoisomer. **(A-C)** Root length was measured from images taken after 8 days using ImageJ. Bars represent mean of normalized root length ± SEM of ≥ 12 biological replicates. Stars indicate significant differences between treatment and 0 μM Aze media *P* < 0.05 in unpaired *t*-test.

To test the effects of altering Pro levels in vivo on Aze tolerance, we isolated two Arabidopsis Pro cycle mutants, which have altered Pro levels. *prodh1-4* (SALK_119334) is a T-DNA insertional mutant in *AtProDH1* (encoding for proline dehydrogenase; At3g30775) the first step of Pro catabolism. This mutant overaccumulates Pro, particularly upon abiotic stress (Funck et al., 2010; Sharma et al., 2011). We also isolated a mutant in Δ1-pyrroline-5-carboxlyate synthetase1 (*AtP5CS1*; p*5cs1-4*; SALK_063517), which catalyzes the first step in Pro biosynthesis from glutamate (Glu) (Verslues and Sharma, 2010) and has reduced Pro levels (Székely et al., 2008; Sharma and Verslues, 2010). Mutants were grown on plates containing 10 μM Aze and compared with wild-type (Wt, Col-0). *prodh1-4* and *p5cs1-4* showed no difference from Wt when grown in the absence of Aze, however *prodh1-4* roots grew longer in the presence of 10 μM Aze compared with Wt (Supplementary Fig. 2). Whereas *p5cs1-4* showed no difference compared with Wt when grown on 10 μM Aze (Supplementary Fig. 2).

We hypothesized that increased Pro levels in *prodh1-4* and decreased Pro in *p5cs1-4* leads to increased and decreased tolerance to Aze, respectively. To further test this hypothesis, we grew mutants on Aze with varying Pro concentrations to determine the exogenous Pro concentration required to fully restore growth. *prodh1-4* consistently showed longer roots compared to Wt at increasing Pro concentrations, whereas *p5cs1-4* consistently showed shorter root lengths compared to Wt (Supplementary Fig. 2). Less exogenous Pro was required to fully restore the root grow of *prodh1-4* compared to *p5cs1-4* (Supplementary Fig. 2). These data indicate that *prodh1-4* is more tolerant to Aze likely because of higher endogenous Pro concentrations.

### Restoration of Aze growth reduction by other amino acids

Next, we tested the effects of the other 19 proteogenic amino acids as well as some Pro metabolic precursors to determine if these could restore Arabidopsis Aze-induced root growth. Surprisingly, some other amino acids restored growth to similar levels as L-Pro when supplied at a 10:1 ratio (100 μM amino acid:10 μM Aze), including the small hydrophobic amino acids Ala and Gly, and the polar amino acid Gln (Fig. 2C). Some amino acids restored growth, but not fully, including the aromatic amino acids Try, Phe, and Trp (Fig. 2C). Whereas most amino acids had no or very little impact on restoring Aze-induced root growth (Fig. 2C). A Pro precursor, ornithine, and a hydroxylated form of Pro were unable to fully restore growth (Supplementary Fig. 3).

To understand how Arabidopsis responds metabolically to Aze treatment, we analyzed free amino acid levels from Arabidopsis grown for 8 days on control media and then moved to 10 μM Aze-containing media for 1, 2, 24, and 48 hours. We expected some amino acids to increase due to the stress imposed by Aze treatment, such as Pro, which is upregulated following abiotic stresses such as drought and salt (Verslues and Sharma, 2010). Individual amino acids showed variable responses (Fig 3, Supplementary Table 1). Many amino acids were relatively unchanged, such as Met, Phe, Glu, and Lys (Fig 3, Supplementary Table 1). Interestingly, Pro levels were unchanged during the Aze treatment (Fig 3, Supplementary Table 1). Some amino acids showed an increase such as Ala, Gly, Ile, and Thr (Fig. 3, Supplementary Table 1). Interestingly, these amino acids were also some of the same amino acids that were able to restore Aze growth reduction (Fig. 2C). These data show that Aze treatment stimulates accumulation of particular amino acids.

**Fig. 3.**
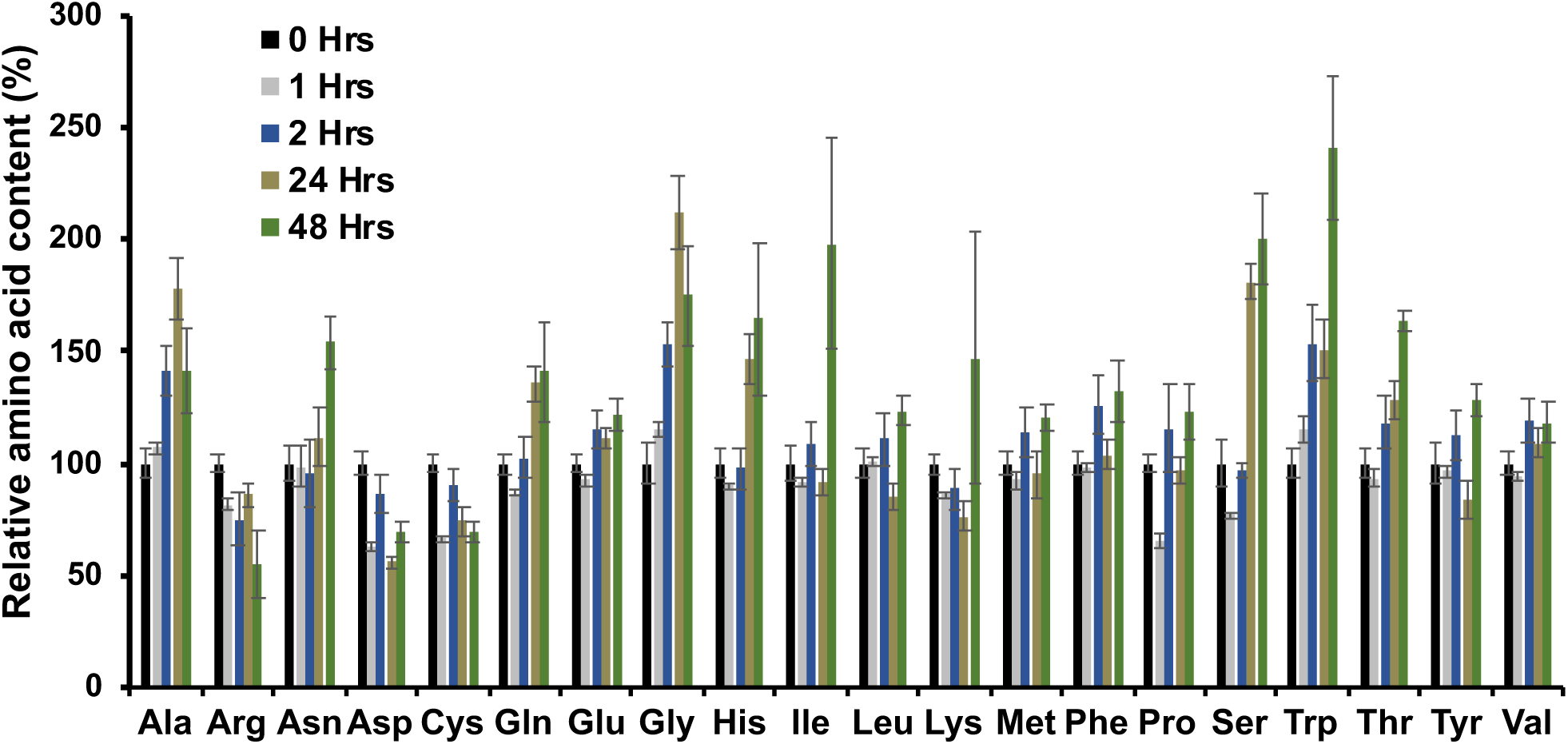
Amino acid response during Aze treatment. Arabidopsis seedlings were grown for 8 days on control media, then moved to 10 μM Aze containing media for 0, 1, 2, 24 and 48 hours. Amino acids were extracted and quantified from total seedlings. Each individual amino acid is normalized to the value at the 0 hour timepoint so that increases and decreases are more obvious. Bars represent normalized mean ± normalized SEM of 4 biological replicates. Full amino acid quantification and statistical analysis can be found in Supplemental Table 1.

### Aze is misincorporated for Pro during protein biosynthesis

Given that L, but not D-Pro fully restores Aze-induced growth reduction (Fig. 2A) we hypothesized that Aze is misincorprated for Pro during protein biosynthesis. To test this hypothesis, we performed untargeted proteomics from Arabidopsis root and leaf tissue grown on 10 μM Aze for 8 days. More than 30,000 confident peptides were identified in all the tissues and treatments, demonstrating the quality and consistency of the protein extractions and proteomics analyses (Fig. 4A). In total around 4,000 unique proteins were detected across the treatments and tissues (Supplementary Tables 2-5). To identify peptides with potential misincorporation events, we treated Aze as a post-translation modification with a mass shift of 14.01 Da compared to Pro during data analysis (Supplementary Fig. 4). Aze was misincorporated in >3.0 % of peptides when grown on 10 μM Aze regardless of tissue (Fig. 4A). Next, we removed peptides that did not contain Pro residues and recalculated the Aze misincorporation rates. From Arabidopsis grown on 10 μM Aze, a total of 20,390 and 16,782 peptides containing Pro residues were detected in the leaves and roots, respectively (Supplementary Tables 3 & 5). This translates into a slightly higher Aze misincorporation rate of 5.9% and 5.6% in the leaves and roots, respectively (Supplementary Tables 3 & 5). We also analyzed the data to identify the total number of Pro residues that were detected and then the number of those residues that had Aze misincorporation. From leaf tissue grown on 10 μM Aze, we detected > 2.5 million Pro residues, 6,491 of those had an Aze misincorporation event, leading to a 2.52 % Aze misincorporation rate (Supplementary Table 3). A similar analysis in the roots identified > 1.9 million Pro residues, 4,881 of those had an Aze misincorporation event, leading to a 2.49 % misincorporation rate (Supplementary Table 5). Regardless of how the proteomics data was analyzed we consistently identify Aze misincorporation, however at relatively low rates.

**Fig. 4.**
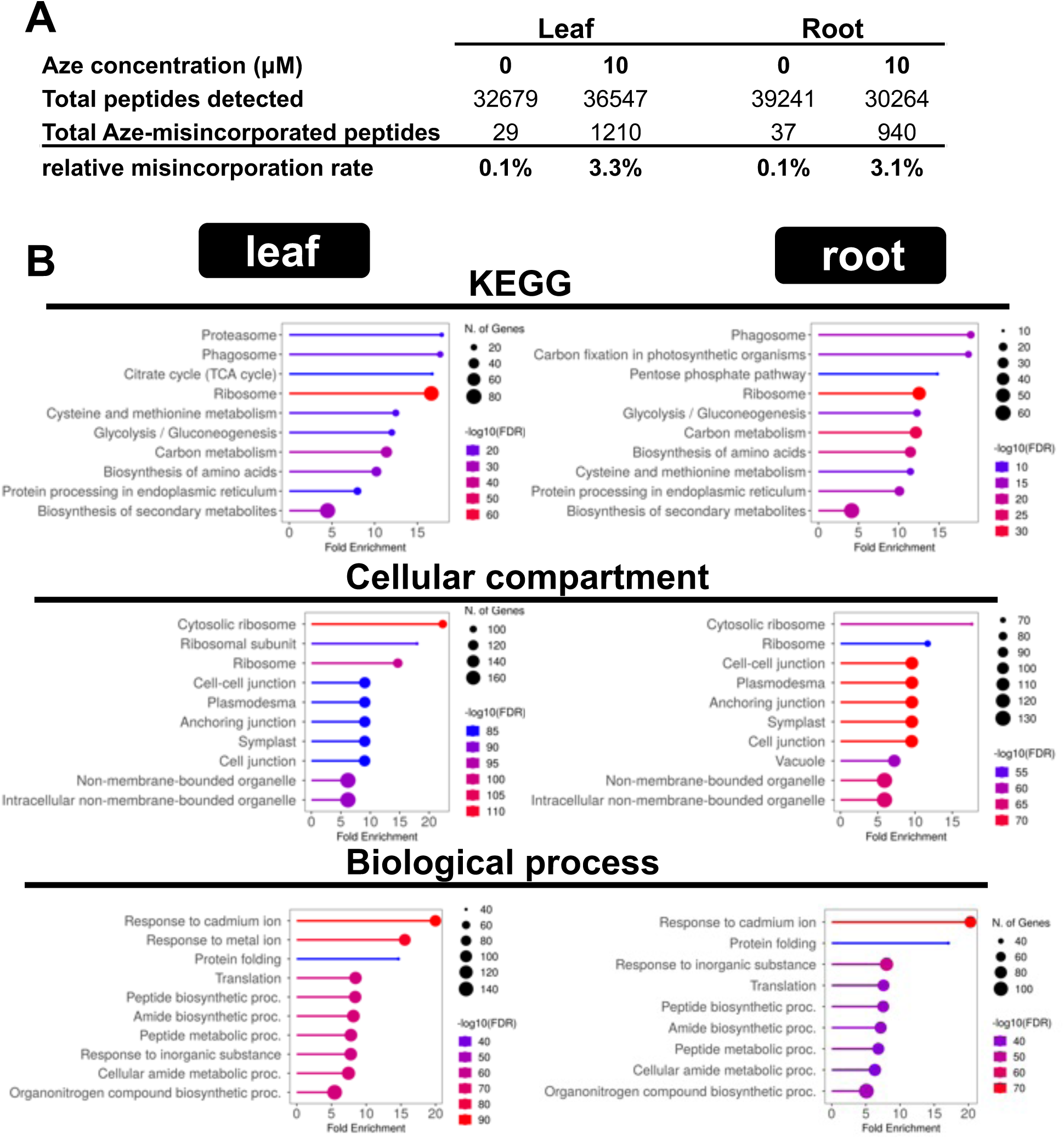
Aze misincorporation in Arabidopsis proteins. **(A)** Total proteins were extracted from leaf and root tissue of Arabidopsis grown on 10 μM Aze for 8 days. Total peptides, and Aze misincorporated peptides are shown for each tissue and Aze concentration. Relative misincorporation rate is the total Aze-misincorporated peptides over the total confidently identified peptides. **(B)** Functional enrichment using ShinyGO 0.80 of proteins with Aze misincorporation in leaf and root using an FDR of 0.05. Size of the circle indicates the number of genes found in that category over the total number of genes in that category, and the color indicates the significance of the enrichment –log (enrichment FDR).

We next analyzed which Pro residues are prone to misincorporation for two abundant proteins that showed Aze misincorporation in the leaves and roots (At5g02500 and At2g36530). At5g02500 contains 26 Pro residues, peptides with Aze misincorporation covered 8 of the 26 Pro residues, which were scattered across the protein (Supplementary Fig 5A). Similarly, At2g36530 contains 16 Pro residues, and 6 were prone to misincorporation. The peptides that matched were distributed across the entire protein (Supplementary Fig. 5B). These data indicate that at least for these two proteins, Aze misincorporation events are distributed across Pro residues throughout the protein sequence and are not preferentially located in certain regions of the protein (Supplementary Fig. 5).

We next analyzed the proteomics data to understand if Aze misincorporation is a random process or if it is enriched in proteins with certain biological functions. Functional enrichment analysis was conducted for those proteins that show Aze misincorporation. Using KEGG pathways, terms such as proteosome, phagosome, ribosome and protein processing at the ER were enriched in both the root and leaf data sets, all indicators of Aze triggering a wider stress response (Fig. 4B). However, other metabolic processes like carbon metabolism and biosynthesis of amino acids were also enriched (Fig. 4B), suggesting that Aze is also affecting central carbon metabolism. Cytosolic and ribosomal cellular compartment terms were enriched, likely indicating the preferential involvement of cytosolic processes (Fig. 4B). The biological processes that were enriched were response to cadmium ion, protein folding, translation and various other terms related to protein biosynthesis and stress (Fig. 4B), further indicating effects on protein biosynthesis and wider stress responses.

Given that Arabidopsis has two Pro-tRNA synthetases with different substrate affinities and subcellular locations (Lee et al., 2016) and that cytosolic processes were enriched in the proteomics data (Fig. 4B) we investigated where Aze misincorporated proteins are synthesized. Proteins translated in subcellular compartments (for example, mitochondria and plastid encoded genes) showed no Aze misincoporation (Supplementary Fig. 6, Supplementary Tables 2-5), whereas nuclear encoded proteins, which are translated in the cytosol or at the ER membrane showed around 3 % Aze misincorporation rate (Supplementary Fig. 6). Some chromosomes had slightly lower Aze misincorporation rates, such as chromosomes 2 and 4, but these lower misincorporation rates are likely due to different chromosomal lengths and less overall genes on those chromosomes. These data indicate that Aze misincorporation affects proteins specifically translated in the cytosol and encoded in the nuclear genome.

### Aze treatment stimulates the unfolded protein response

The unfolded protein response (UPR) is an ER stress response due to accumulation of misfolded proteins (Howell 2012). Given that Aze is misincorporated for Pro in Arabidopsis proteins, and that ER processes and the proteosome were enriched in the proteomics analysis, we hypothesized that Aze misincorporation triggers a wider stress response. To test this hypothesis, we treated Arabidopsis with Aze to determine if the UPR response was triggered. As a control, we used tunicamycin, a known UPR stimulator that interferes with glycosylation of proteins processed in the ER resulting in accumulation of unfolded proteins (Chen and Brandizzi, 2013; Howell, 2013). Following treatment with Aze or tunicamycin for 8 days, qRT-PCR was performed targeting two genes involved in the UPR response (binding protein 1,2 (BiP1,2) and BiP3; Liu et al., 2007; Iwata et al., 2008; Liu and Howell, 2010). As expected *AtBiP1,2* (*At5g28540, At5g42020*) and *AtBiP3* (*At1g09080*) gene expression was induced after 8 days of tunicamycin treatment (Fig. 5A). Plants grown on 10 μM Aze had an increased expression of *AtBiP3,* but to a lesser degree than treatment with tunicamycin (Fig. 5A). *AtBiP3* was induced by 123-fold following tunicamycin treatment and 18-fold following Aze treatment (Fig 5A). Aze treatment for shorter times (2 and 4 hours) was not sufficient to stimulate the UPR (Supplementary Fig. 7). Amino acid profiling during a time course of Aze treatment shows that the total pool of free amino acids increased by about 40% during the Aze treatment (Fig. 5B). These data indicate that increase in free amino acids may be due to protein turnover and further indicate that a wider stress response is triggered following Aze treatment involving the UPR.

**Fig. 5.**
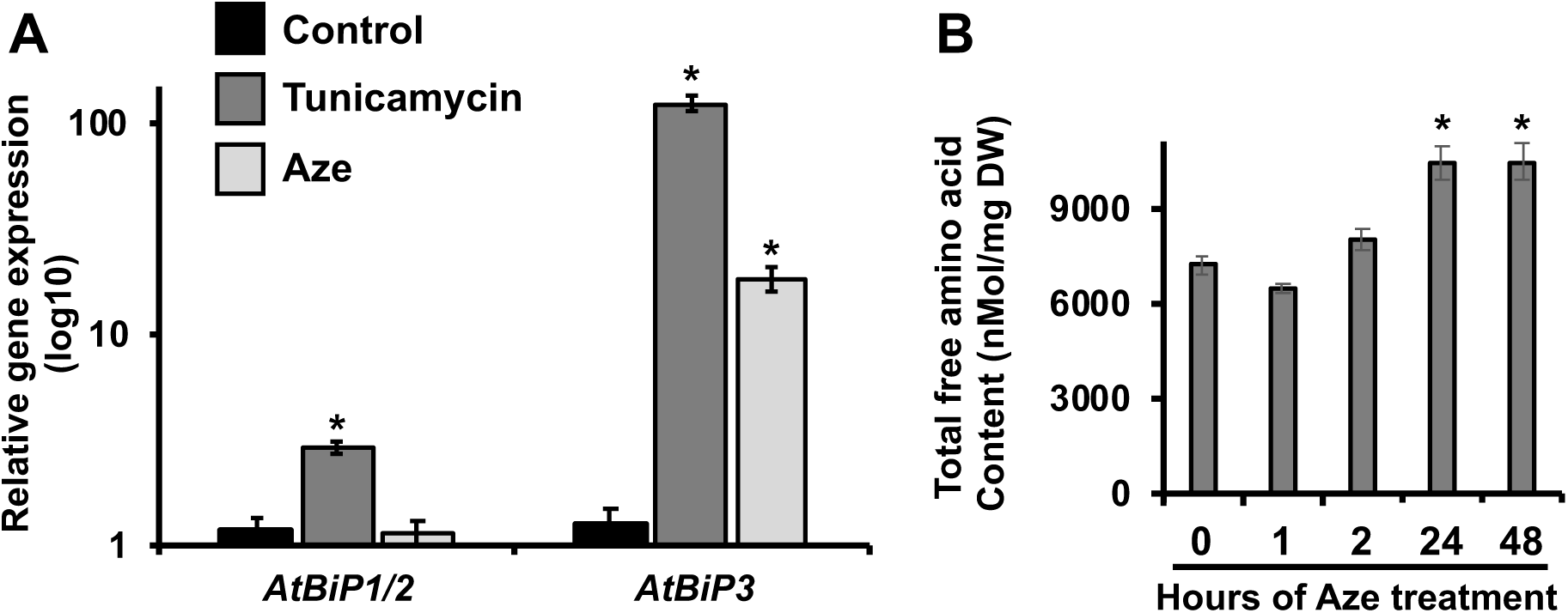
Aze triggers the unfolded protein response. **(A)** Gene expression analysis using qRT-PCR was performed on RNA extracted following growth on 10 μM Aze or 10nM tunicamycin for 8 days. UPR response was monitored following gene expression of *AtBiP1/2* and *AtBiP3*. Bars indicate mean ± SEM of 4 > biological replicates. Stars indicate significant differences between treatment and control *P* < 0.05 in unpaired *t*-test. **(B)** Amino acids were extracted from whole seedlings grown on control media for 8 days and then moved to Aze for 1, 2, 24, and 48 hours. Total amino acids are the sum of all 20 individually detected free amino acids. Bars indicate mean ± SEM of 4 biological replicates. Stars indicate significant differences between treatment and 0 hr time point *P* < 0.05 in unpaired *t*-test.

## Discussion

Structurally diverse metabolites are key mediators in plant-environment interactions and enable adaptation to harsh environments. The varied structures of plant specialized metabolites translate into diverse biological activities and modes of actions. However, the lack of knowledge of modes of action limits our ability to co-opt plant metabolites for agricultural, industrial, or medicinal purposes. Here, we identified the mode of action of Aze in plants providing insight into ways to engineer tolerance to Aze and other biologically active metabolites.

### Aze is broadly toxic, targeting a conserved molecular mechanism

Phylogenetically diverse plants all showed reduced root growth on Aze-containing media (Supplementary Fig. 1). However, the potency of Aze varied depending on the plant. Some organisms show enhanced tolerance to Aze, for example two different microbes have evolved enzymes that can detoxify Aze (Nomura et al., 2003; Gross et al., 2008). A strain of the green alga *Nannochloris bacillaris* was isolated that accumulates 26-fold more Pro than wild-type leading to Aze tolerance (Vanlerberghe and Brown, 1987). Additionally, Aze-producing plants have evolved resistance strategies, such a highly specific Pro-tRNA synthetase found in *D. regia* that can exclude Aze (Norris and Fowden, 1972). This contrasts with the Arabidopsis cytosolic Pro-tRNA synthetase, which binds Aze and Pro equally (Duchêne et al., 2005; Lee et al., 2016), and is likely the reason why Arabidopsis and other organisms are susceptible to Aze. These studies highlight that Aze is a broadly toxic metabolite and that there is natural variation in the types of resistance mechanisms, which could be used to engineer Aze tolerance into crops.

### Growth restoration of hydrophobic amino acids

Aze growth reduction could be restored by adding L-, but not D-Pro (Fig. 2A). Further, proteomics shows that Aze was misincorporated for Pro during protein biosynthesis (Fig. 4). Surprisingly, some small hydrophobic amino acids, Ala, Gly, had restorative effects when complementing Aze growth reduction (Fig. 2C). One possible scenario is that there is competition for import between these amino acids and Aze from non-specific amino acid importers in the roots (Lehmann et al., 2011; Tegeder, 2012). Given that there was a 10:1 ratio of amino acid:Aze in this experiment, the hydrophobic amino acids may be preferentially imported over Aze in the roots, thus no or very little growth reduction is observed (Fig. 2C). Another reason Ala may rescue growth could be competition between Aze and Ala for binding Pro-tRNA-synthetases. Pro, Aze, and Ala can bind to *E. coli* and Hunan Pro-tRNA synthetases (Beuning et al., 2000). However, Ala, unlike Aze and Pro, is hydrolyzed from the tRNA synthetase prior to misincorporation (Beuning et al., 2000). Arabidopsis could have the same proofreading mechanism on their Pro-tRNA synthetases and the competition between Ala and Aze for the Pro-tRNA synthetase active site might explain why adding Ala rescues Aze-induced root inhibition.

### Aze misincorporation leads to a global stress response

Given the relatively strong root growth reduction phenotype observed when plants are grown on Aze (Fig. 1, Supplementary Fig. 1), yet the relatively low Aze misincorporation rate, we hypothesized that Aze may be triggering a more global stress response. Indeed, the UPR response was upregulated following 8 days of Aze treatment (Fig. 5A). These data suggest that Aze misincorporation leads to protein misfolding. Misfolded proteins accumulate in the cell and trigger the UPR response at the ER and likely other locations triggering a wider stress response. Functional enrichment of proteins with Aze misincorporation events also indicates a global response (Fig. 4B). AtBiP3 – an ER chaperone induced during the UPR - was detected in our proteomics data and showed Aze-misincorporation in both the leaves and roots (Supplementary Tables 2-5). Furthermore, the amino acid profiling suggests that Aze treatment triggers protein degradation, as total pools of free amino acids increase during Aze treatment (Fig. 5B). Other biologically active plant metabolites with different structures can have a similar effect on perturbing protein folding and function (Zer et al., 2020; Meyer et al., 2021) and suggest plants have independently evolved structurally distinct metabolites to interfere with protein biosynthesis/function.

### Aze misincorporation specifically on nuclear encoded genes

We observed a dramatic difference in Aze-misincorporation for nuclear encoded proteins compared to proteins encoded in the plastid or mitochondria genome. Nuclear encoded genes that would be translated on ribosomes in the cytosol or associated with the ER and targeted to their subcellular compartment show around a 3% Aze misincorporation rate (Supplementary Fig. 6). Whereas genes encoded in the plastids or mitochondria show no Aze misincorporation (Supplementary Fig. 6). One explanation could be that Aze is not imported into the plastids or mitochondria and not available for protein synthesis. This seems unlikely as Pro can be imported and incorporated in protein synthesis in the plastids and mitochondria. Arabidopsis has two Pro-tRNA synthetases and they are localized in different locations, one to the cytosol and one with dual localization to the plastids and mitochondria (Duchêne et al., 2005). The cytosolic Pro-tRNA synthetase binds Aze and Pro equally, while the plastid and mitochondrial form is about 100-fold more specific for Pro than Aze (Lee et al., 2016). The high specificity of the organellar Pro-tRNA synthetase likely explains why no misincorporation is observed for proteins synthesized in the plastids or mitochondria. It is likely that plants that make Aze have similarly specific Pro-tRNA synthetases (Norris and Fowden, 1972). Structural insight into the differences in plant Pro-tRNA synthetases will provide information about key residues and motifs that enable specificity.

Identification of the mode of action of Aze in plants enables greater understanding of how plants that make Aze can tolerate its toxicity. Plants that accumulate Aze are found primarily in two distinct orders (Asparagales and Fabales; Fowden, 1956; Fowden and Steward, 1957; Sung and Fowden, 1971; Leete et al., 1974). Although the precise distribution within these orders is not known, the most parsimonious explanation is that Aze biosynthesis arose through convergent evolution, as has been observed for other plant metabolic pathways such as caffeine (Pichersky and Lewinsohn, 2011; Huang et al., 2016; Agrawal, 2017; Meyer et al., 2021).

Further phylogenetic placement of Aze accumulation across plants will enable greater understanding of the evolution of the Aze pathway (Strauss and Agrawal, 1999; Schenck and Busta, 2021). Additional studies on the plants that make Aze will provide insight into whether tolerance mechanisms also evolved through convergent evolutionary mechanisms and provide tools to engineer plants to produce biologically active metabolites for enhanced resilience.

## Materials and methods

### Plant growth conditions

Arabidopsis seedlings (Col-0 ecotype) were grown on 0.6% agar-solidified medium with 0.5x Murashige and Skoog Basal Medium with Vitamins (PhytoTech Labs, Lenexa, KS, USA) and 1% sucrose. The media was supplemented with filter-sterilized (.22 μm) L-azetidine-2-carboxylic acid (Aze, Sigma) or other amino acids of varying concentrations as defined in the figure legends. Seeds were surface sterilized with chlorine gas (80 mL bleach + 20 mL 12 M hydrochloric acid) for 1 hour in a desiccator. Sterilized seeds were placed on solidified media in square petri dishes (100 mm x100 mm) and plates were oriented vertically. Prior to position plates under lights, they were placed at 4°C for 48 hours. Growth conditions were the following: ∼23°C, 16 h daylight and 8 h dark photoperiod and light intensity was ∼ 6800 lux. Arabidopsis was grown for 8 days, unless noted otherwise in the figure legends, photographed, and root lengths were measured using ImageJ (Schneider et al., 2012). Other plants grown on Aze including *Psychine stylosa*, *Solanum lycopersicum* (tomato Beefsteak), *Nicotiana tabacum* (K326), *N. benthamiana*, and *Lactuca sativa* (lettuce, Black Seeded Simpson variety were sterilized using 90 % ethanol (v/v) for 1 minute, followed by 15 % bleach (v/v) for 15 minutes, and 5 washes with autoclaved H_2_0 for 6 minutes each. Growth times varied depending on the plant and time to germination.

### IC*_50_* Calculations of Arabidopsis growth on Aze

Growth conditions were the same as previously described for Arabidopsis, with a gradient concentration of Aze (2.5 μM; 5 μM; 7.5 μM; 10 μM; 25 μM; 50 μM; 100 μM). IC_50_ values were calculated from normalized root lengths of ≥ 8 biological replicates per concentration using a four-parameter logistic regression model (Sebaugh, 2011).

### Arabidopsis growth recovery following Aze treatment

Arabidopsis was grown on 0.6% agar-solidified medium with 0.5x Murashige and Skoog Basal Medium with Vitamins (PhytoTech Labs, Lenexa, KS, USA) and 1% sucrose containing 10 μM Aze for 4 or 8 days. Seedlings were transferred in a laminar flow hood to new media containing either 0 or 10 μM Aze. Root lengths were measured as previously described after 12 days.

### Isolation of Arabidopsis Pro cycle T-DNA mutants

*prodh1-4* (SALK_119334) and *p5cs1-4* (SALK_063517) were ordered from TAIR. Homozygous lines were identified by PCR-based genotyping using the primers listed in (Supplementary Table 6). DNA was extracted from 4-week-old plants using newly emerged young leaf tissue (∼2mm square size) added to microfuge tubes with 600 μL of lysis buffer (lysis buffer: 10 mM Tris-HCl pH 8.0, 25 mM ethylenediaminetetraacetic acid (EDTA), 0.5 % (w/v) sodium dodecyl sulfate (SDS)). The leaf tissue was ground with lysis buffer then incubated at 55 °C for 15 minutes. 200 μL of 5 M ammonium acetate was added and vortexed for 20 seconds. The samples were centrifuged at 14,000g for 3 minutes and 600 μL of the supernatant was transferred to a separate tube. An equal volume (∼600 μL) of isopropanol was added and centrifuged at 14,000g for one minute and the supernatant was removed. The pellet was rinsed with 400 µL of 70 % (v/v) ethanol and spun at 14,000g for 1 minute. The supernatant was removed, and the pellet was dried in a laminar flow hood for 30 minutes. The pellet was resuspended in 50 µL water.

Template DNA was used with gene and T-DNA specific primers (Supplementary Table 6) and Phusion DNA polymerase (Thermo) were mixed. The following PCR programs were used: *AtP5CS1* initial denaturation at 98 °C for 30 sec, then 30 cycles of denaturation at 98 °C for 10 sec, annealing at 60 °C for 15 sec, extension at 72 °C for 1 min, final extension at 72 °C for 10 min. *AtProDH1* initial denaturation at 98 °C for 30 sec, then 30 cycles of denaturation at 98 °C for 10sec, annealing at 60 °C for 20 sec, extension at 72 °C for 1 min, final extension at 72 °C for 10 min. Homozygous lines were identified (Supplementary Fig. 8) and used for plate-based growth assays as described above.

### Proteomics analysis of Aze misincorporation

Arabidopsis was grown on 10 μM Aze for 8 days and proteins were extracted from roots and leaves and similarly for plants grown on control media without Aze. Three biological replicates were performed consisting of ∼10 plants each separated into root and leaf tissue. Tissue was ground to a fine powder in liquid nitrogen and proteins were extracted in 100 mM Tris pH 7.8 with 5% (w/v) SDS and 10 mM dithiothreitol (DTT). The extraction was heated at 65 °C for 20 minutes and centrifuged at 16,000g for 20 minutes. Proteins were alkylated with 30 mM iodoacetamide, precipitated using methanol and chloroform precipitation and washed once with 80% (v/v) cold acetone. Protein (50 μg) were digested with LyC at 1:50 (enzyme:protein) for 3 hours at 37 °C and then digested with trypsin (1:50, enzyme:protein ratio) overnight at 37 °C and purified by Evosep tips.

For all proteome analyses, we used an EvoSep One liquid chromatography system (Bache et al., 2018). A 15 cm × 150 μm internal diameter column with 1.5 μm C18 beads (Bruker PepSep) and a 10 µm internal diameter zero dead volume electrospray emitter (Bruker Daltonik). Evosep One was coupled online to a modified trapped ion mobility spectrometry quadrupole time-of-flight mass spectrometer (timsTOF Pro 2, Bruker Daltonik GmbH, Germany) via a nanoelectrospray ion source (Captive spray, Bruker Daltonik GmbH).

Data were analyzed by timsTOF Pro2 in DDA PASEF mode. PASEF and TIMS were set to “on”. One MS and ten PASEF frames were acquired per cycle of 1.17sec (∼1MS and 120 MS/MS). Target MS intensity for MS was set at 10,000 counts/sec with a minimum threshold of 2500 counts/s. Duty cycle was locked to 100%. Ion mobility coefficient (1/K0) value was set from 0.6 to1.6 V.s/cm^2^, collision energy was set from 20-59 eV. MS data were collected over m/z range of 100 to 1700. If the precursor (within mass width error of 0.015 m/z) was >4X signal intensity in subsequent scans, a second MS/MS spectrum was collected. Exclusion was active after 0.4min. Isolation width was set to 2 for m/z <700 or 3 for m/z>700.

PEAKS version 10.6 was used to search the data against Arabidopsis TAIR10 database appended with reversed protein sequences as decoys. For the data analysis, precursor and fragment mass tolerances were set to 20 ppm, and 0.1 Da. Up to two missed protease cleavages were allowed. Oxidation of methionine and Pro to Aze (delta mass =-14.0156 Da) were set as variable modifications. Carbamidomethylation of Cysteine was set as a fixed modification. Maximum number of variable modifications per peptide was set to 3. Data were exported from PEAKS and peptides and proteins were filtered to FDR ≤0.01, Ascore >=20. Functional enrichment analysis was performed on proteins with Aze misincorporation using ShinyGO 0.80 (Ge et al., 2020). Size of the circle indicates the number of genes found in that category, and the color indicates the significance of the enrichment (Fig. 4).

### Localization of proteins with Aze-misincorporation

Nuclear encoded genes were assumed to be translated on cytosolic ribosomes or on ribosomes associated with the ER. Involvement of cytosolic tRNA synthetases would be involved in these processes. Nuclear encoded gene were identified by the At locus ID, those with Atxg where the x through 1-5 indicated nuclear encoded on one of the five chromosomes. Plastid and mitochondrial encoded proteins were identified by their At locus ID with AtCg or AtMg.

### Amino acid quantification during Aze treatment

Arabidopsis seedlings were grown for 8 days on media without Aze, then transferred to 10 μM Aze-containing media for 0, 1, 2, 24, and 48 hours. Seedlings were flash frozen in liquid nitrogen, four biological replicates were performed containing at least 7 seedlings each. Seedlings were lyophilized for 48 hours and pulverized with 3 mm glass beads using a Spex GenoGrinder. Free amino acids were extracted, detected, and quantified by liquid chromatography-tandem mass spectrometry (LC-MS/MS) as previously described (Yobi et al., 2020; Thomas et al., 2024).

### Quantification of unfolded protein response (UPR)

The UPR response was quantified from Arabidopsis seedlings grown for 8 days on 10 μM Aze-containing media and in seedlings that were grown for 8 days on media without Aze, then transferred to 10 μM Aze-containing media for 2 and 4 hours. As a positive control, tunicamycin – a known stimulator of the UPR response (Ruberti and Brandizzi, 2018) - was used at 10 nM at the same time points as Aze. Following treatments, seedlings were flash frozen in liquid nitrogen, pooled, and ground to a fine powder using a mini pestle. RNA was extracted using a Qiagen RNaeasy extraction kit and cDNA was synthesized using All-In-One 5X RT Master Mix (Applied Biological Materials) containing DNase I. Gene expression of *AtBiP1,2* (*At5g28540, At5g42020*) and *AtBiP3* (*At1g09080*) (Liu et al., 2007; Iwata et al., 2008; Liu and Howell, 2010), were monitored using qPCR. cDNA was diluted five times and additional five-fold dilutions were made to calculate primer efficiency. All primer pair efficiencies were between 90%–100%. cDNA was mixed with master mix containing 5 μL of Fast SYBR Green Master Mix (Applied Biosystems) and 200 nM of each primer. Primer sequences used for qRT-PCR can be found in Supplementary Table 6. Reactions were placed in a Quantstudio3 Real-Time PCR system (Applied Biosystems) using the following PCR cycle: an initial denaturation at 95 °C for 1 min, 40 cycles of amplification at 95 °C for 3 sec, 60 °C for 30 sec. Ct values were extracted for each reaction and used to quantify initial cDNA concentration using 2^−ΔΔC(t)^ method normalized to a control gene (*At2g28390*; Hong et al., 2010) for relative quantification.

## Author Contributions

WTS conducted experiments, analyzed data, and revised the manuscript. VD conducted experiments, analyzed data, and revised the manuscript. HND conducted experiments, analyzed data, and revised the manuscript. MM conducted experiments, analyzed data, and revised the manuscript. KVH conducted experiments, analyzed data, and revised the manuscript. RA guided and oversaw the project, and revised the manuscript. CAS planned, guided and oversaw the project, performed experiments, analyzed data, wrote and revised the manuscript.

## Acknowledgements

We thank Adam Yokom for valuable insight and feedback on this study. We thank Brian Mooney and Thao Nguyen from the Charles W. Gehrke Proteomics Facility at MU for assistance in proteomics analyses. We acknowledge support from MU and a College of Agriculture, Food and Natural Resources Joy of Discovery Grant (CAS).

## Supplementary Information

**Supplementary Fig. 1.**
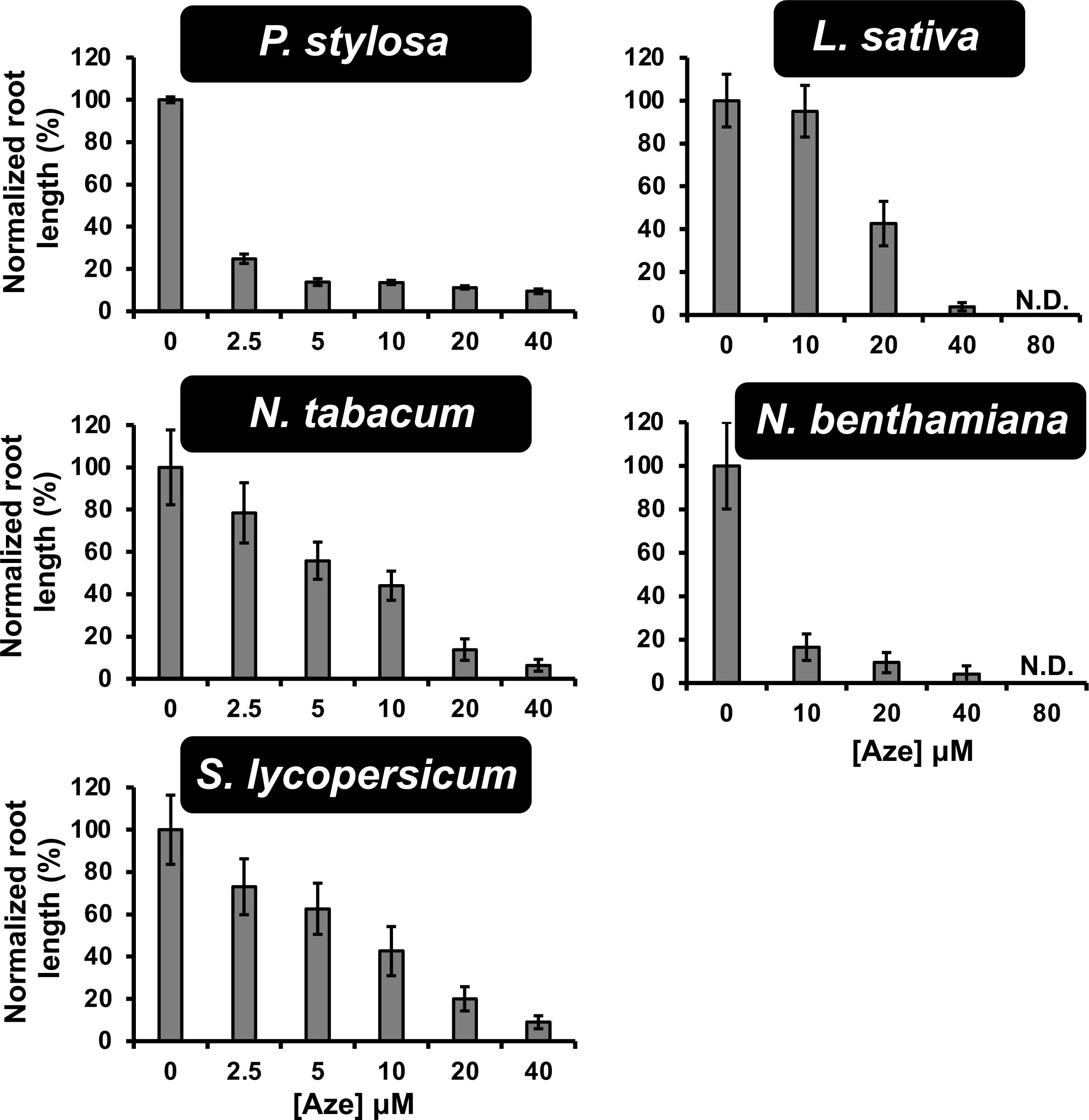
Diverse plants are susceptible to Aze. Plants were sterilized and germinated on media containing increasing concentrations of Aze. Images were taken and root lengths were calculated by tracing roots in ImageJ. Bars represent normalized mean ± normalized SEM of ≥ 6 biological replicates. N.D. No root length detected.

**Supplementary Fig. 2.**
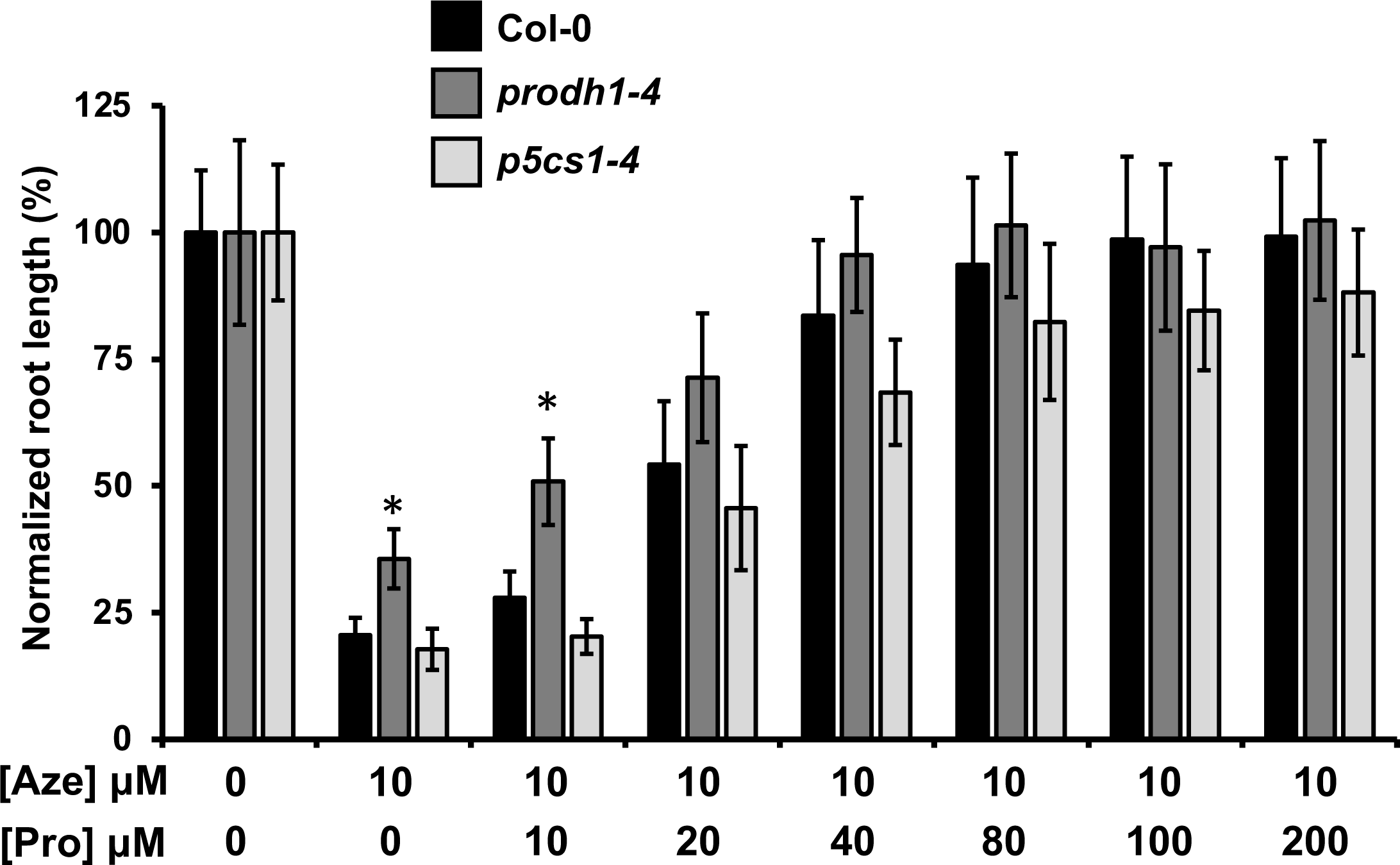
Arabidopsis Pro cycle mutants show altered response to Aze and Pro. Arabidopsis wild-type (Col-0) and two Pro cycle mutants, *prodh1-4* and *p5cs1-4*, which have enhanced and decreased Pro levels following abiotic stress, respectively were grown on 10 μM Aze with increasing Pro supplemented in the media from 10-200 μM. Bars indicate normalized mean ± normalized SEM of ≥ 6 biological replicates, stars indicate significant differences between mutant and Col-0 at each concentration with *P* < 0.05 in unpaired *t*-test.

**Supplementary Fig. 3.**
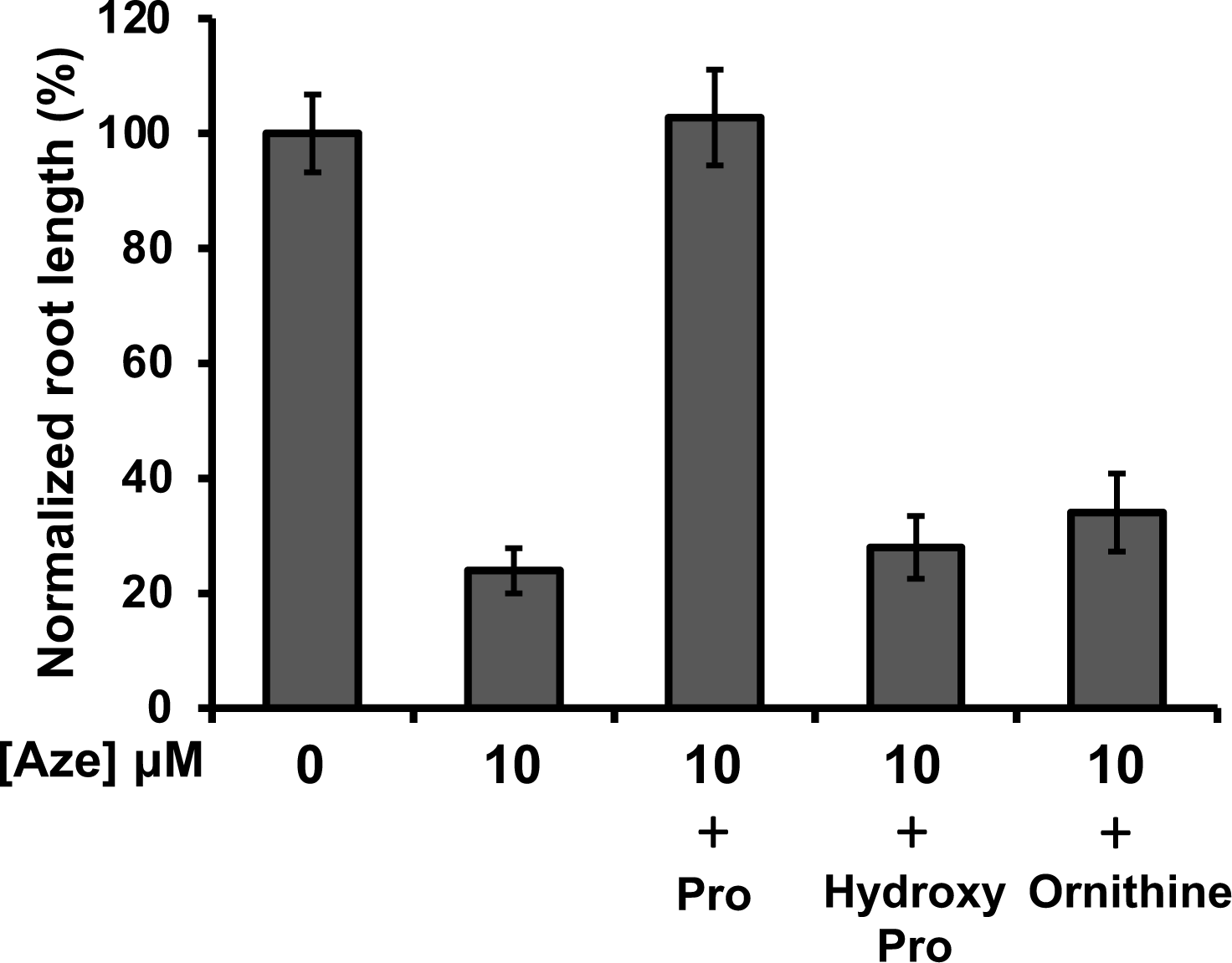
Arabidopsis response to growth on Aze supplemented with other metabolites. Arabidopsis was grown on 10 μM Aze and supplemented with 100 μM of L-Pro, trans-4-hydroxy-L-Pro, or L-ornithine. Bars indicate normalized mean to the 0 μM Aze control ± normalized SEM of ≥ 8 biological replicates.

**Supplementary Fig. 4.**
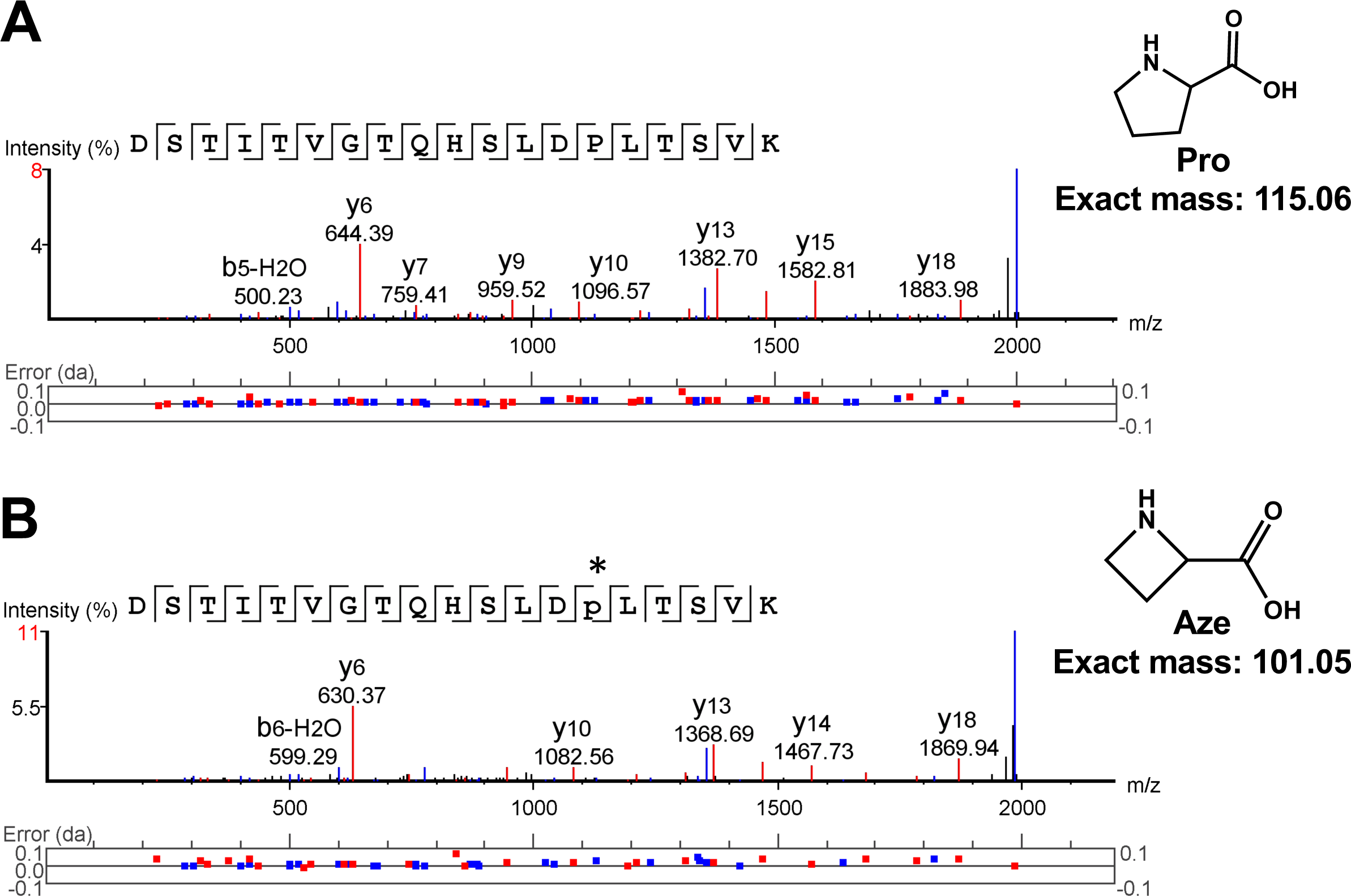
Example MS/MS spectra of peptide with Aze misincorporation. MS/MS spectra showing a series of b (blue) and y-ions (red) for a peptide that was detected with no Aze incorporation **(A)** and with Aze incorporation **(B)**. Aze misincorporation was identified by searching for post translational modifications (PTM) on Pro of a mass difference of -14.0156 Da. The Pro residue with Aze misincorporation is shown with a star. Structures of Pro and Aze are shown.

**Supplementary Fig. 5.**
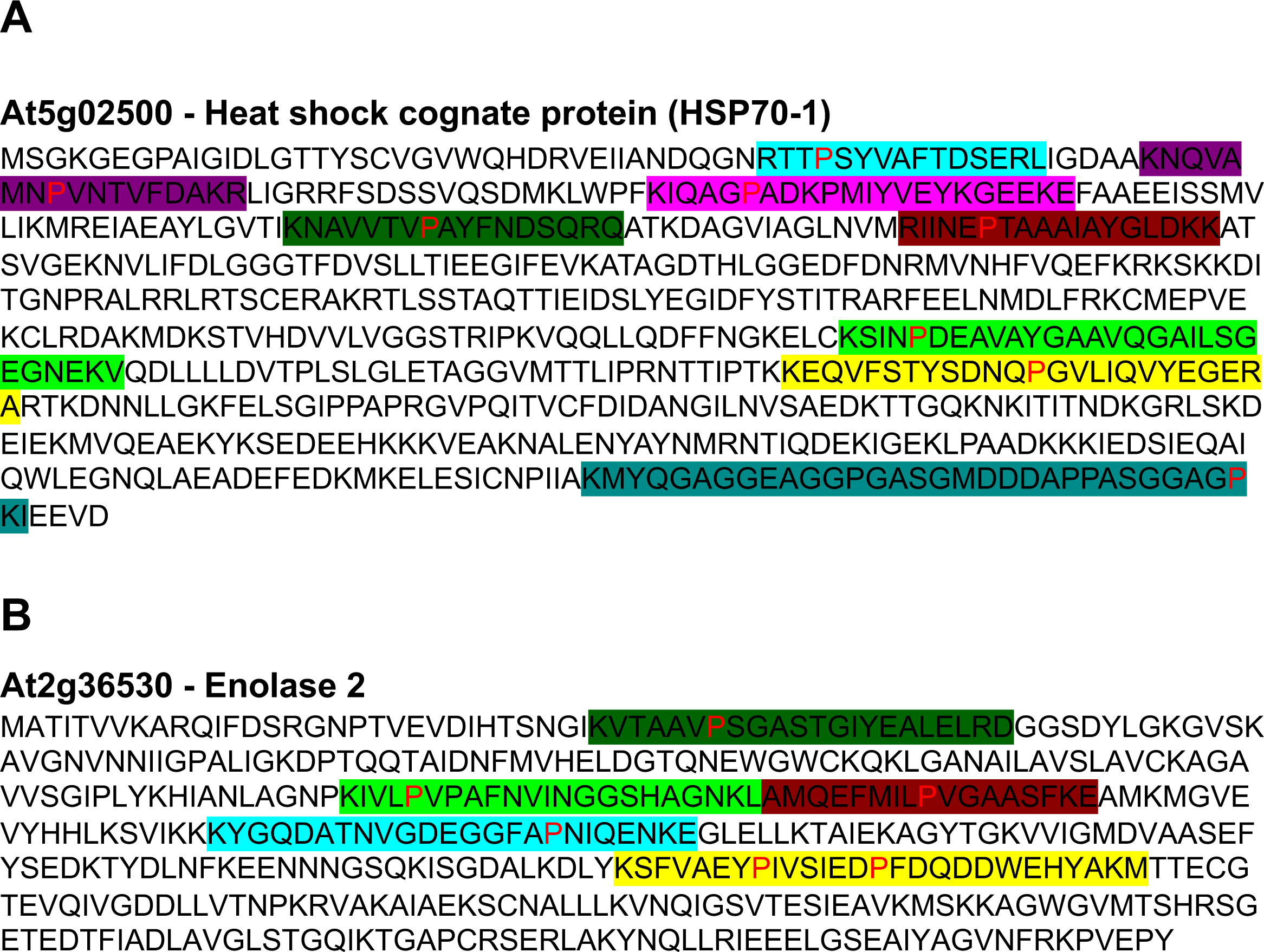
Distribution of detected peptides with Aze misincorporation across two proteins. Sequences of two proteins are shown and the highlighted regions indicates detected peptides that match the protein sequence. Red highlights indicate the Pro residue that had an Aze misincorporation. These two proteins were high abundant proteins that also showed high Aze miscincorporation in multiple peptides. Aze misincorporation events are found distributed across the whole protein sequence **(A)** At5g02500 encodes a heat shock cognate protein. **(B)** At2g36530 encodes an enolase 2 protein.

**Supplementary Fig. 6.**
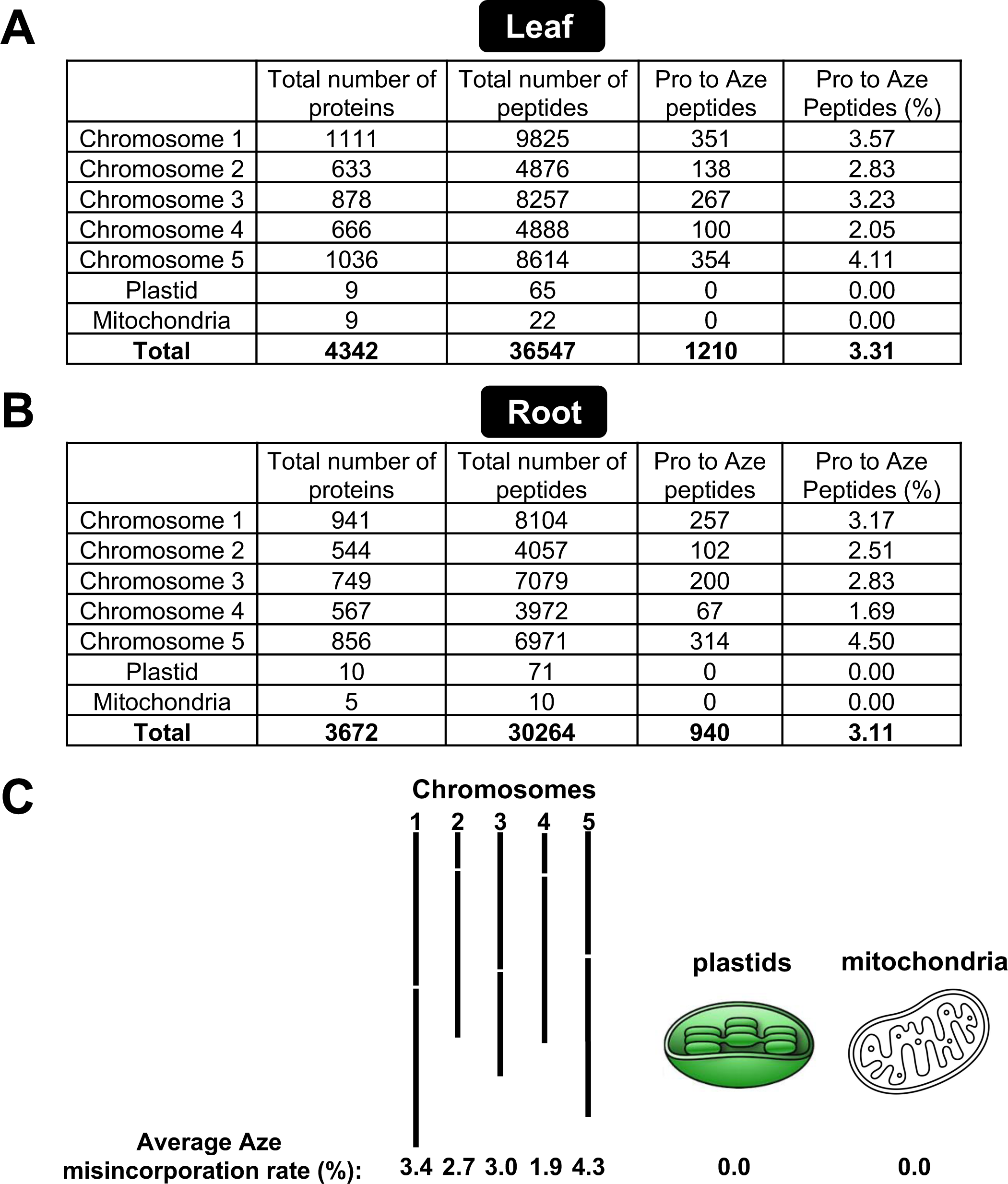
Aze misincorporation is specific to nuclear encoded proteins. **(A)** Aze misincorporated peptides from leaf tissue grown on μM Aze grouped by which genome they are located on, nuclear, plastid or mitochondrial. If nuclear encoded, they were separated based on chromosome location. **(B)** Aze misincorporated peptides from root tissue grown on 10 μM Aze grouped by which genome they are located on, nuclear, plastid or mitochondrial. If nuclear encoded, they were separated based on chromosome location. **(C)** Average of root and leaf **(A** and **B)** mapped onto cartoon diagrams of chromosomes and plastid and mitochondrial genomes. Full proteomics data sets can be found in Supplementary Tables 2-5.

**Supplementary Fig. 7.**
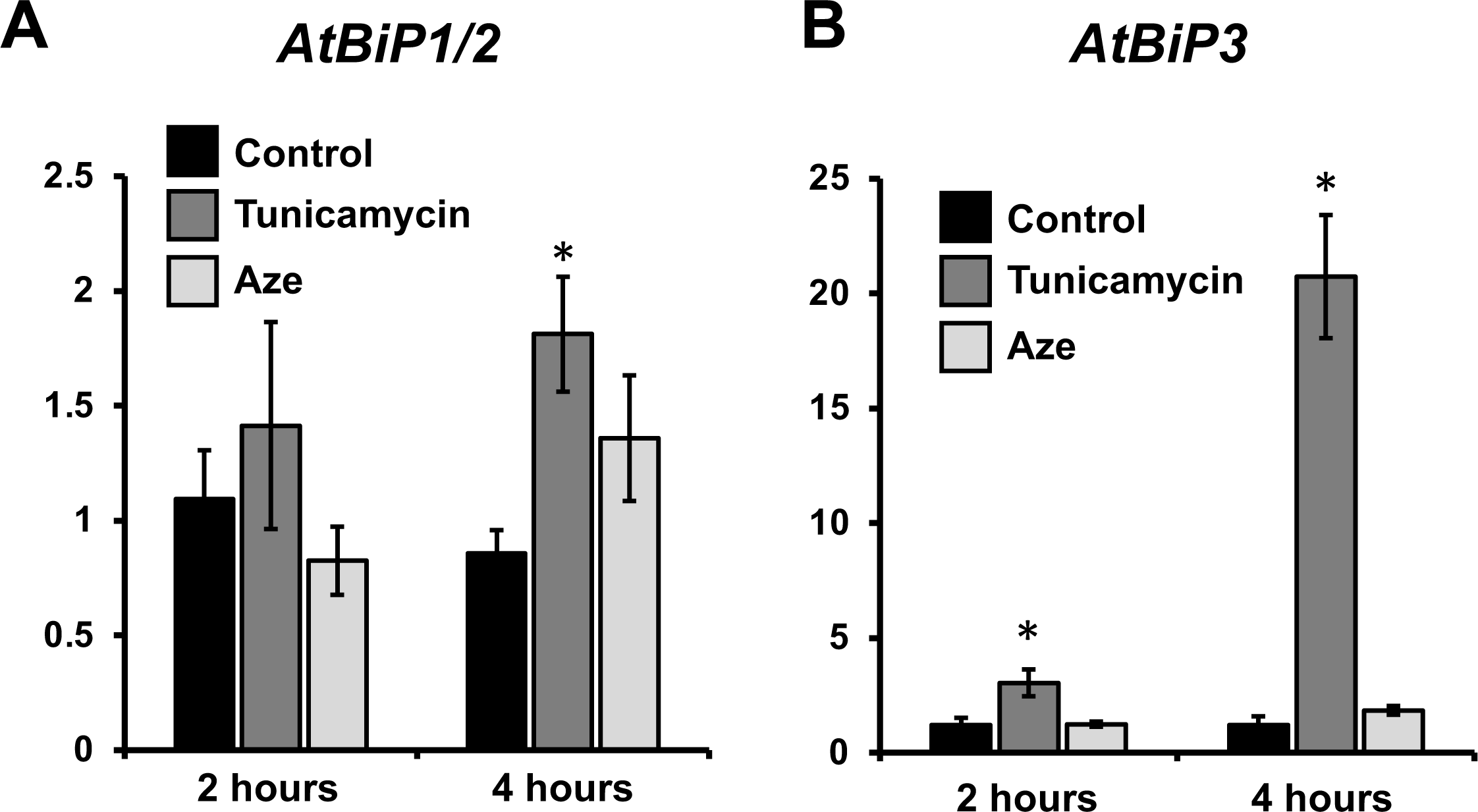
UPR response genes are not activated after 2 and 4 hours of Aze treatment. **(A)** Gene expression analysis using qRT-PCR was performed on RNA extracted following 2 and 4 hours of 10 μM Aze, 10 nM tunicamycin treatment or control. UPR response was monitored following gene expression of **(A)** *AtBiP1/2* and **(B)** *AtBiP3*. Bars indicate mean ± SEM of 4 > biological replicates. Stars indicate significant differences between treatment and control with *P* < 0.05 in unpaired *t*-test.

**Supplementary Fig. 8.**
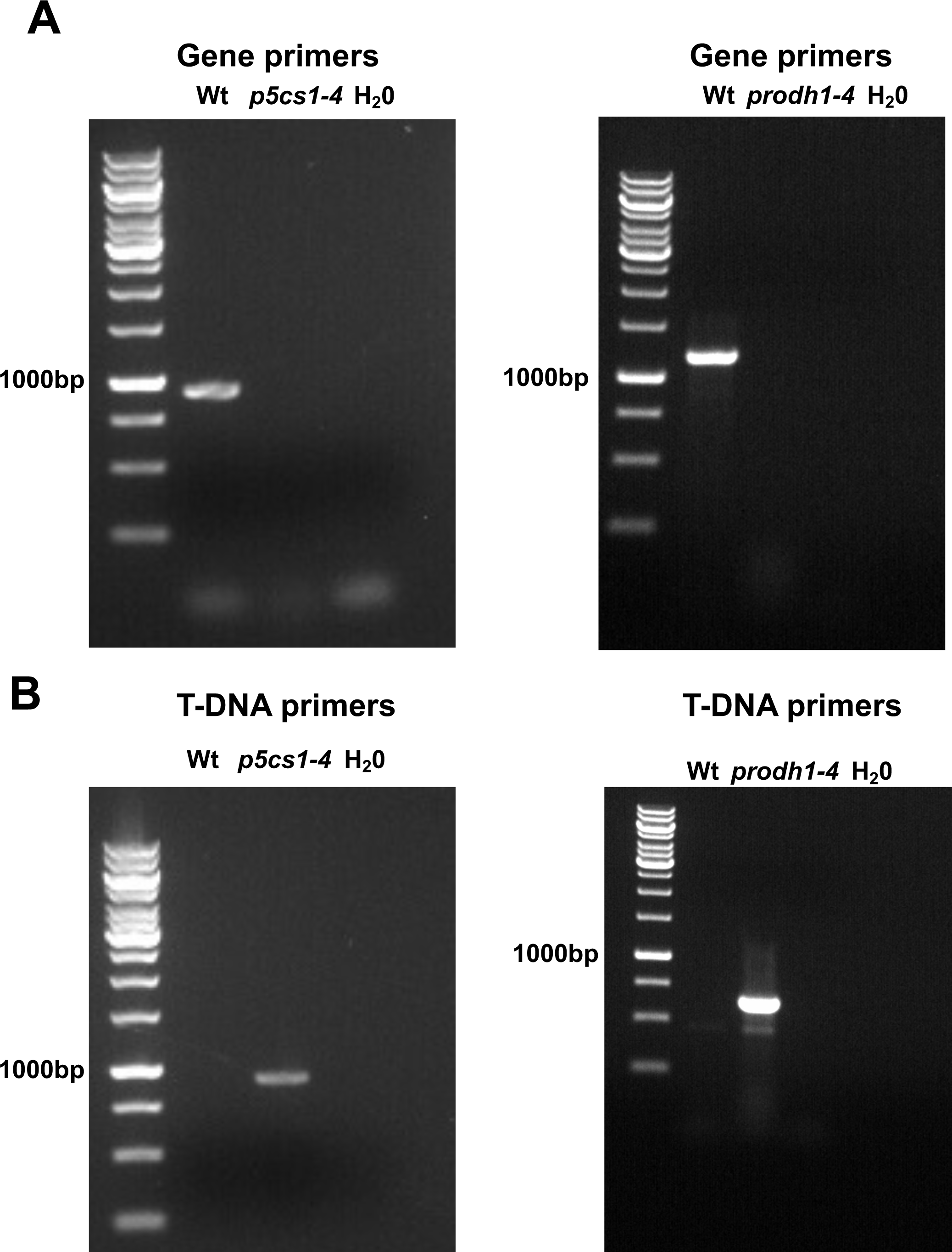
Genotyping of Arabidopsis Pro cycle T-DNA mutants. Homozygous *p5cs1-4* and *prodh1-4* mutants were identified using PCR with gene-specific primers **(A)** and a gene-specific with a T-DNA specific primer **(B)**. Homozygous lines were identified by a single band produced in the T-DNA PCR **(B)** and no amplification with gene specific primers **(A)**. Primers used are listed in Supplementary Table 6.

